# Potent neutralization antibodies induced by a recombinant trimeric Spike protein vaccine candidate containing PIKA adjuvant for COVID-19

**DOI:** 10.1101/2021.02.17.431647

**Authors:** Jiao Tong, Chenxi Zhu, Hanyu Lai, Chunchao Feng, Dapeng Zhou

**Affiliations:** Tongji University School of Medicine, Shanghai, 200092, China

## Abstract

Neutralizing antibodies are critical to prevent corona virus infection. The structures of immunogens to elicit most potent neutralization antibodies are still under investigation. Here we tested the immunogenicity of the trimeric, full length Spike protein with 2 proline mutations to preserve its prefusion conformation. Recombinant trimeric Spike protein expressed by CHO cells was used with polyI:C (PIKA) adjuvant to immunize mice by 0-7-14 day schedule. The results showed that Spike-specific antibody was induced at day 21 with titer of more than 50,000 in average as measured by direct binding to Spike protein. The titer of neutralization reached more than 1000 in average when tested by a pseudo-virus system, using monoclonal antibodies (40592-MM57 and 40591-MM43) with neutralizing IC50 at 1 μg/ml as standards. Protein/peptide array showed that the antibodies induced by trimeric S protein vaccine bind similarly to natural infection with the receptor binding domain (RBD) as major immunodominant region. No linear epitopes were found in RBD, although several linear epitopes were found in C-terminal domain right after RBD, and heptad repeat regions. Our study supports the efficacy of recombinant trimeric Spike protein vaccine candidate for COVID-19, with excellent safety and readiness for storage and distribution in developing countries.

## Introduction

It is generally accepted that effective COVID-19 vaccines are the only approach to end the global pandemic. Currently, inactivated virus (1), Adenoviral vector-based vaccines (2-4) and mRNA vaccines (5-6) encoding Spike protein have been approved for urgent use in China, US, and other countries. However, these vaccines do not meet the need for vaccination in all countries. For example, the inactivated COVID-19 vaccines are limited by manufacturing capacity due to difficulties in producing live viruses. The currently approved mRNA vaccines require cold-chain transport by freezers at -80 °C or -20 °C, although thermostable mRNA vaccine candidate has been invented (7). The other unanswered question is the duration of immune responses induced by above vaccines.

Vaccines based on recombinant proteins and adjuvants have been approved for HBV, HPV, and Influenza. The manufacturing of recombinant proteins and adjuvants are easy to scale-up, that provides unlimited supply of immunogens. Such vaccines have been proven to be safe and effective. More importantly, the immune responses often last for years. Thus vaccines with the recombinant Spike protein and adjuvants as components may be among best choices for developing countries to end the COVID-19 pandemic.

Both full-length Spike protein and engineered subunit of Spike protein containing receptor binding domain have been reported as vaccine candidates (8-18). In this study, we report a trimeric full length Spike protein containing PIKA (polyI:C) adjuvant. We chose such composition for several reasons: 1) The full length Spike protein contains more T cell epitopes that are essential for inducing viral-specific T cells; 2) The conformation of the trimeric form of Spike protein is similar to S-trimer structure in natural virions; 3) The PIKA (polyI:C) adjuvant has shown excellent safety and efficacy in rabies vaccines by 0-7-14 vaccination schedule (19).

## Results

### Trimeric S protein induced higher neutralizing antibodies

The titer of antibody binding to Spike trimer protein was above 50,000 in average after 3 immunizations by trimeric Spike protein with polyI:C adjuvant. Using a pseudo-virus system, we determined the neutralizing titer to be higher than 1000 in average. Two monoclonal antibodies (40592-MM57 and 40591-MM43) with known neutralizing activities were used as standards for neutralization assays (with IC50 at 1 μg/ml). With same dose of adjuvant and proteins, trimeric Spike protein induced significantly higher neutralizing antibodies than monomeric Spike protein (Figure 1).

**Figure 1.**
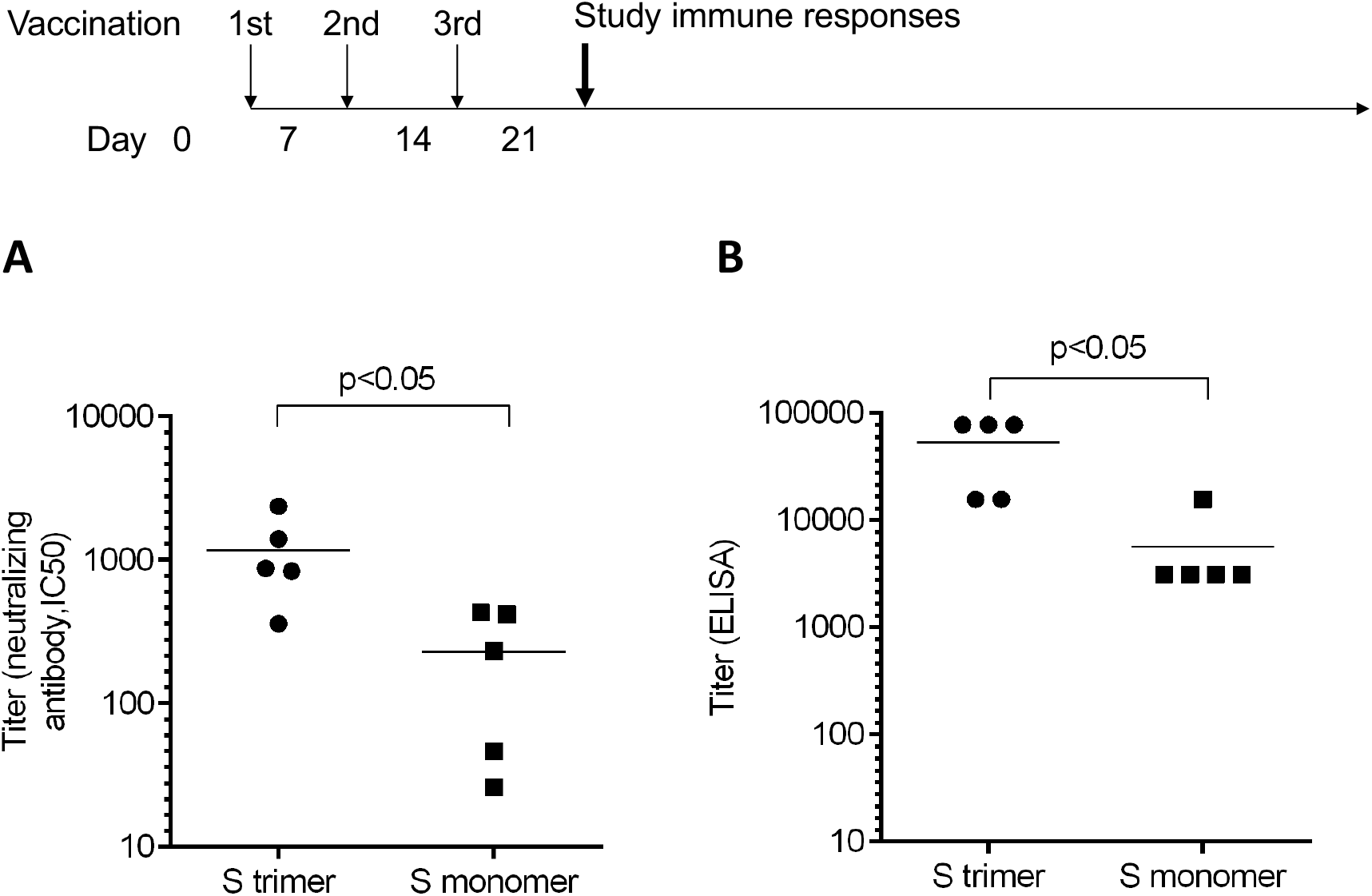
Antibody titers measured by neutralizing assays and ELISA. Mice were immunized by Spike trimer or monomer proteins containing PIKA adjuvant by a 0-7-14 day schedule. A. The neutralizing titer as measured by pseudo-virus with monoclonal antibodies 40592-MM57 and 40591-MM43 as control (with IC50 at 1 μg/ml). B. Antibody titer as measured by ELISA using plate-bound trimeric Spike protein.

### Trimeric S protein induced similar epitope patterns as natural infection

To understand the epitopes of antibodies induced by trimeric Spike protein vaccine, we performed protein/peptide array containing recombinant RBD, S1, and linear peptides of Spike protein (20). Serum from mice vaccinated by trimeric Spike protein vaccine showed strongest binding to RBD, S1 subunit, and S proteins. However, linear epitopes were only observed in the C-terminal domain right after RBD, and heptad repeat regions (Figure 2). Few linear epitopes were found for RBD region, indicating that the observed antibody binding to RBD region are non-linear confirmational epitopes. These results are highly consistent with the epitope patterns of serum from patients with natural infection of COVID-19 (21-22). Our data support the hypothesis that the trimeric Spike protein induced antibody responses similar to natural virions.

**Figure 2.**
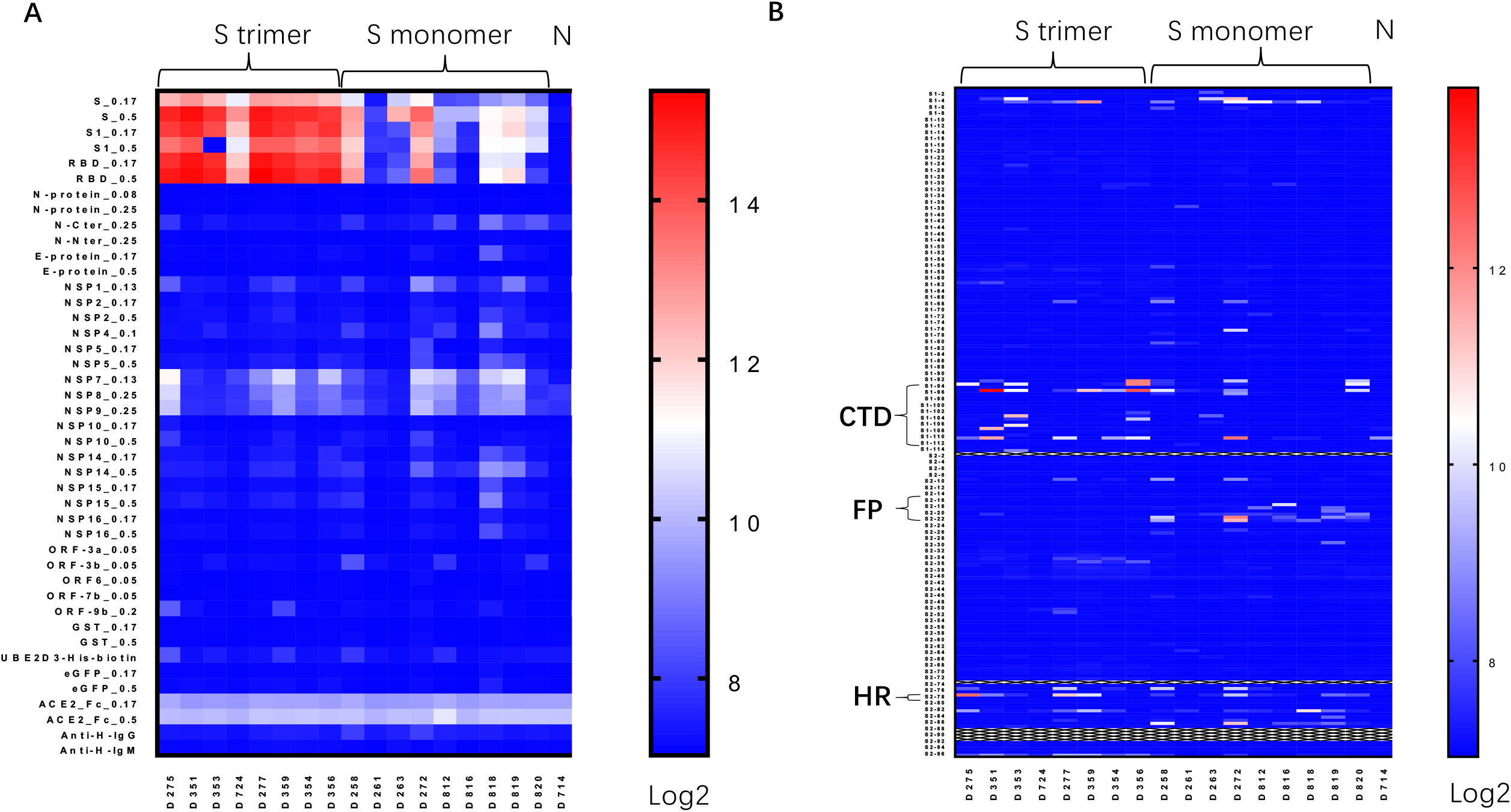
Protein/peptide array of serum antibodies induced by Spike protein vaccines. A. Protein array assay for sera from mice immunized by Spike trimer, Spike monomer, Non-immunized mice, using Spike (S_0.17 and S_0.5 means proteins were printed at 0.17 or 0.5 mg/ml); S1 subunit of Spike, RBD and other viral proteins. B. Linear peptide array using linear peptides of Spike proteins. CTD, C-terminal domain right after RBD (Peptides S1-93-S1-113); FP, fusion peptide (Peptides S2-14-S2-23); HR, heptad regions (Peptides S2-78).

## Discussion

### The neutralizing versus binding ratio of vaccine induced antibodies

The recombinant trimeric Spike protein with polyI:C adjuvant induced neutralizing antibodies with the titer higher than 1000 in average. The comparison to antibody titers induced by mRNA vaccine, inactivated virus and adenoviral vectors remain to be studied. According to previous studies by Walsh et al. (23), the ratio of neutralizing/binding ratio of antibodies induced by BTN vaccine is about 10 fold lower than antibodies from natural infected patients (1:7 by natural infection as compared to 1:20 to 1:40 by vaccination). The mechanisms for lower neutralizing/binding ratio might be due to the incomplete translation, and/or incorrect folding of mRNA encoding Spike protein. The neutralizing/binding ratio of antibodies induced by trimeric Spike protein in human individuals will be evaluated in near future when human subjects are vaccinated.

### The epitope map of antibody induced by vaccines versus natural infection

The similar antibody binding epitopes identified by protein/peptide array suggest that the trimeric recombinant vaccine’s confirmation is similar to Spike protein of natural virions. Noteworthy, monoclonal antibodies with neutralizing activities have been identified to recognize N-termianl domain (NTD, 24). Ma et al identified neutralizing antibodies which bind to C-terminal domain right after RBD (CTD) and fusion peptide (FP) region (21). Structural analysis by Cryo-EM and molecular modeling indicated that heptad repeat (HR) regions are surface-exposed and serve as targets for neutralizing antibody binding (25-27). Clearly, vaccines that can target the non-RBD region play important role in preventing the pandemic of mutant viruses with RBD mutations that escape the RBD-focused vaccines.

## Materials and methods

### Expression and purification of the recombinant SARS-CoV-2 Spike protein trimer and spike protein monomer

To express the prefusion S ectodomain, a gene encoding residues 1−1208 of 2019-nCoV S (GenBank: MN908947) with proline substitutions at residues 986 and 987, a “GSAS” substitution at the furin cleavage site (residues 682–685), a C-terminal T4 fibritin trimerization motif, and an 8XHisTag was synthesized and cloned into the pcDNA3.1 vector. The plasmid was transfected to 293T cells and the recombinant S protein trimers were purified by Ni-NTA (nickel-nitrilotriacetic acid) chromatography (QIAGEN, Germany), followed by size exclusion to further purify the trimers. The purified trimers were verified by SDS-PAGE analysis under non-reducing conditions.

### Vaccination in mice

Animal experiments were approved by the institutional board of Tongji University School of Medicine. Purified recombinant SARS-CoV-2 S-trimer-His tagged protein, or Spike protein monomer (25) was resuspended in PBS (pH 7.4) with PIKA adjuvant (19, TLR3 agonist provided by Yisheng Biopharma Ltd, China). C57BL/6 mice at 6 to 8 weeks old were immunized by intramuscular injection of PIKA-S-trimer vaccine or PIKA-S-monomer vaccine. Every mouse was immunized by 5 μg of S-trimer or S-monomer protein and 50 μg of PIKA adjuvant. Mice were immunized on Day 0, Day 7 and Day 14. Sera were collected on Day 21.

### S-protein specific antibody determined by ELISA

To measure the Spike-specific antibody, 96-well microplate (NUNC-Immuno, Thermo, Waltham, MA) was coated by 50 μl 1 μg/ml S-trimer in PBS (pH 7.4) at 37°C for one hour, washed five times by 0.05% tween in PBS (PBS/T) on a mini-shaker. The plates were blocked by 1% bovine serum albumin (Sigma, St Louis, MO) in PBS at 37°C for one hour and washed by PBS/T for five times. Sera were diluted 25 times in the first well followed by five-fold serial dilution for ten times. Sera were incubated at 37 °C with plate-bound S protein for one hour and washed with PBS/T for five times. Then goat anti-mouse IgG conjugated HRP (Southern Biotech, Birmingham, Alabama) was added with 5000 fold dilution in PBS, and incubated at 37 °C for one hour. After washing for five times, chromogenic substrates were added and incubated for half an hour. The reaction was stopped with H_2_SO_4_ solution (1M). The absorbance was measured at 450 nm and the antibody titer was calculated with GraphPad Prism 7.0 (San Diego, CA).

### Protein/peptide array

The protein/peptide array was performed as described (21). Briefly, peptide-BSA conjugates as well as S protein, S1 protein, RBD protein, and other protein of SARS-CoV-2, were printed in triplicate on PATH substrate slide (Grace Bio-Labs, Oregon, USA) to generate identical arrays in a 1 × 7 subarray format using Super Marathon printer (Arrayjet, UK). The microarrays were stored at -80°C until use. The arrays stored at -80°C were warmed to room temperature and then incubated in blocking buffer (3% BSA in PBS buffer with 0.1% Tween 20) for 3 h. A total of 400 μL of diluted sera or antibodies was incubated with each subarray for 2 h. The arrays were washed with PBST and bound antibodies were detected by incubating with Cy3-conjugated goat anti-mouse IgG (Jackson ImmunoResearch, PA, USA), which were diluted for 1: 1,000 in PBST. The incubation was carried out at room temperature for 1 h. The microarrays were then washed with 1×PBST and dried by centrifugation at room temperature and scanned by LuxScan 10K-A (CapitalBio Corporation, Beijing, China) with the parameters set as 95% laser power/ PMT 480. The fluorescent intensity was extracted by GenePix Pro 6.0 software (Molecular Devices, CA, USA).

### SARS-CoV-2 pseudo-virus production

The Extracellular domain of SARS-CoV-2 spike protein of that (GenBank: MN908947) was engineered in a pcDNA3CMV-based-plasmid as Zhou-COVID-19-Spike (Plasmid #161029, Addgene) to assemble pseudo-virus more efficiently. Portions of VSV-G for production of pseudo-virus was used to replace the signal peptide and transmembrane region. The plasmids of 9μg pHAGE-luciferase-GFP、 psPAX2 and Zhou-COVID-19-Spike were co-transfected into HEK293T cells by using 1 μg/mL polyetherimide (Polysciences, Warrington, PA) in DMEM medium containing 10% FCS. After 48 and 72 hours, the supernatant was harvested and pooled. The supernatant containing pseudo-virus was centrifuged at 3000g and filtered through a 0.45μm sterilized membrane (Millipore, Burlington, MA). The titer of virus generated by engineered Zhou-COVID-19-Spike plasmid is ten-fold higher than non-engineered Spike protein sequence (data not shown), and remained stable after two rounds of freeze-thawing. The virus was stored in -80 °C as culture supernatant and used directly for antibody neutralization assays without further purification.

### Neutralization of serum antibody against pseudovirus infection

A 293T-ACE2 cell line (293T/ACE2) expressing human ACE2 was used for virus neutralization assay. 3×10^4^ cells per well were seeded on 96 well plates 12 hours before infection. 50 μL pseudo-virus was incubated with equal volume of serially diluted antibodies for one hour at 37 °C. Monoclonal antibodies 40592-MM57 and 40591-MM43 (Sinobiological, Beijing, China) were tested as standards in parallel, in the concentrations ranging from 0.1 μg/mL to 100 μg/mL. The mixtures of pseudo-viruses and antibodies were added to 293T/ACE2 cells. After 12 hour co-incubation, the co-culture medium was replaced with fresh DMEM containing 10% fetal bovine serum, and the samples were incubated for an additional 48 hours at 37 °C. Luciferase substrate (Promega, Madison, WI) was added at 100 μL per well to lyse the cells. The fluorescence was read by a microplate reader (TECAN). The 50% neutralization dose was calculated using GraphPad Prism 7.0 (San Diego, CA).

## Acknowledgement

We thank Dr. Ming-liang Ma and Prof Sheng-ce Tao (Shanghai Jiao Tong University) for help on protein and peptide array analysis. This work was supported by National Key Research and Development Plan grant 2017YFA0505901, Fundamental Research Funds for the Central Universities 22120200163, National Natural Science Foundation of China grant 31870972, Shanghai Science and Technology Commission grant 15002360172,the Outstanding Clinical Discipline Project of Shanghai Pudong (PWYgy2018–10).

